# Selective Removal of 7KC by a Novel Atherosclerosis Therapeutic Candidate Reverts Foam Cells to a Macrophage-like Phenotype

**DOI:** 10.1101/2023.10.23.563623

**Authors:** Prerna Bhargava, Darren Dinh, Fadzai Teramayi, Ana Silberg, Noa Petler, Amelia M. Anderson, Daniel M. Clemens, Matthew S. O’Connor

## Abstract

The removal of the toxic oxidized cholesterol, 7-ketocholesterol (7KC), from cells through the administration of therapeutics has the potential to treat atherosclerosis and various other pathologies. While cholesterol is a necessary building block for homeostasis, oxidation of cholesterol can lead to the formation of toxic oxysterols involved in various pathologies, the most prominent of which is 7KC, which is formed through the non-enzymatic oxidation of cholesterol. Oxidized LDL (oxLDL) particles, highly implicated in heart disease, contain high levels of 7KC, and molecular 7KC is implicated in the pathogenesis of numerous diseases, including multiple sclerosis, hypercholesterolemia, sickle cell anemia, and multiple age related diseases. Of particular interest is the role of 7KC in the progression of atherosclerosis, with several studies associating elevated levels of 7KC with the etiology of the disease or in the transition of macrophages to foam cells. This research aims to elucidate the molecular mechanisms of UDP-003, a novel therapeutic, in mitigating the harmful effects of 7KC in mouse and human monocyte and macrophage cell lines. Experimental evidence demonstrates that administration of UDP-003 can reverse the foam cell phenotype, rejuvenating these cells by returning phagocytic function and decreasing both reactive oxygen species (ROS) and intracellular lipid droplet accumulation. Furthermore, our data suggests that the targeted removal of 7KC from foam cells with UDP-003 can potentially prevent and reverse atherosclerotic plaque formation. UDP-003 has the potential to be the first disease-modifying therapeutic approach to treating atherosclerotic disease.

## 1. Introduction

Atherosclerosis is a chronic inflammatory cardiovascular disease that is the leading cause of global mortality (Zhu et al. (2018)). It is characterized by the accumulation of lesions within arterial blood vessel walls comprised of lipid deposits, smooth muscle cells, fibrous proliferations, foam cells, and other detritus (Zhu et al. (2018)). Lipid deposits infiltrating the arterial wall are composed of low-density lipoproteins (LDL) that can be modified through oxidation to form oxLDL (Chistiakov et al. (2018); Nilsson et al. (2007)). Progression of atherosclerosis causes the narrowing of blood vessels and results in the rupture of the atherosclerotic plaque, which can cause an array of cardiovascular events including stroke, heart failure, and myocardial infarction (Frostegård (2013)). Accumulation of LDL, oxLDL, and 7KC stresses the arterial wall, thereby activating proinflammatory cytokines in endothelial cells (Chistiakov et al. (2018)). OxLDL contains many oxysterols, but the most abundant oxysterol is 7KC, a toxic and highly proinflammatory molecule that causes significant damage to membranes, pathways, and overall cellular function (Anderson et al. (2019); Rao et al. (2014); Vejux et al. (2020)). 7KC is a byproduct of the non-enzymatic oxidation of cholesterol, most commonly found in oxLDL or around inflamed cells and tissues. Prior studies show significantly elevated 7KC in the blood cells of CVD patients and within atherosclerotic plaque (Tang et al. (2018)). 7KC initiates an inflammatory response in endothelial cells (EC), which facilitates recruitment of immune cells, including monocytes, which differentiate into macrophages. These differentiated macrophages engulf oxidized forms of cholesterol, including 7KC. Due to subtle but important structural differences between 7KC and cholesterol, intake of excessive 7KC leads to the transformation of macrophages into foam cells, one of the primary components of atherosclerotic plaque (Calle et al. (2019); Pirillo et al. (2013)). Through dysfunction of their lysosomal system, foam cells become engorged with lipids and are incapable of participating in reverse cholesterol transport (RCT). Foam cells subsequently start emitting stress signals, recruiting more (doomed) macrophages, and eventually erupt and die, releasing anything they may have consumed as well as their cellular components into the plaque, making it even bigger. In vivo studies have further demonstrated cellular dysfunction in macrophages prompted by 7KC through inflammation (Chang et al. (2021); Terasaka et al. (2008)).

Oxidative stress plays a pivital role in the development of atherogenesis. Oxidative stresses entails reactive oxygen and nitrogen species (ROS, RNS). Under normal physiological conditions, ROS and RNS serve as signaling molecules essential for maintaining fundamental cellular functions. An array of intrinsic antioxidative defense systems eliminates RNS and ROS in cells once they have performed their signaling functions. However, unregulated production of ROS leads to cellular damage, as the presence of ROS oxidizes cellular protein, lipids, and DNA and ultimately causes cellular death. ROS is implicated in the progression of atherosclerosis, as it both perpetuates chronic inflammation within the vascular wall and participates in the formation of foam cells. According to the oxidative modification hypothesis, when LDL is modified through oxidation, this leads to alteration in apolipoprotein B, rendering LDL particles recognizable by scavenger receptors on macrophages. This leads to further uptake of cholesterol in macrophages, resulting in the formation of foam cells (Steinberg et al. (1989)).

One of the main characteristics of foam cells is an accumulation of intra-cellular lipid droplets, which give the cells a bloated and “foamy” appearance. (Brown et al. (1979)). RAW 264.7 murine macrophages have been shown to accumulate lipid droplets and transform into foam cells under exposure to 7KC (Calle et al. (2019)). It has been reported that both dysfunction in RCT and an increase in oxLDL/7KC uptake contribute to lipid droplet accumulation. Under normal conditions, RCT shuttles cholesterol out of cells, allowing HDL particles to pick up the esterified cholesterol to be taken to the liver and either recycled or excreted. However, cholesterol efflux from foam cells becomes greatly reduced, serving as the rate-limiting step for RCT. Cholesterol efflux is modulated by ATP binding cassette (ABCA1) expression. ABCA1 is crucial for effluxing and transporting cholesterol to the liver via RCT (Attie et al. (2001)). ABCA1 is regulated by liver X receptor alpha (LXR*α*) (Tian et al. (2009); Venkateswaran et al. (2000)). When macrophages become foam cells, the expression of ABCA1 protein and the LXR receptor decreases, making it more difficult for the cells to remove free cholesterol (Zhao et al. (2014)). PPAR*γ*, a nuclear receptor, is pivotal in regulating the ATP binding cassette and several studies have shown that oxLDL activates PPAR*γ* (Nagy et al. (1998); Ricote et al. (1998); Taketa et al. (2008)).

A second distinctive feature of foam cells is decreased phagocytic ability. Phagocytosis is an essential process by which macrophages, the sentinel of immune cells, internalize apoptotic cells, pathogens, and cellular debris in vacuoles, where they are recycled or degraded (Hirayama et al. (2017)). During disease progression, 7KC presence leads to increased oxidative stresses, intracellular lipid accumulation, and inability to phagocytose, leading to an increase in apoptosis in macrophages (Calle et al. (2019); Palozza et al. (2010)). Apoptosis is highly regulated programmed cell death mediated through ROS, oxidative stress, and dysfunction in the lysosome and endoplasmic reticulum, leading to the accumulation of lipids in the cytoplasm of cells (Su et al. (2019)). Necrosis is an irreversible process of cellular death where the cell membrane ruptures, leading to additional detrimental effects on the surrounding environment. During atherosclerotic progression, both these processes play an essential role.

Multiple clinical trials have reported elevated 7KC levels in cardiovascular disease patients (Song et al. (2017); Wang et al. (2017)). Lipidomic analysis in patients with heart failure has also revealed a significant elevation of 7KC in erythrocytes and plasma compared to healthy individuals (Tang et al. (2018)). These results indicate that 7KC is a promising therapeutic target in ameliorating diseases. Cyclodextrins (CDs) are cyclic glucose oligomers with a hydrophobic cavity and a hydrophilic shell. This unique shape allows them to spontaneously interact with many different hydrophobic guest molecules and form a water soluble, noncovalent CD host-guest inclusion complex (Fig. 1) (Loftsson et al. (2004)). To help improve their solubility and/or specificity towards one guest molecule over another, functional groups such as methyl (Me), sulfobutyl (SB), and hydroxypropyl (HP) groups can substitute the hydroxyl groups on the glucopyranoside subunits of the native CD. Some CDs, such as HP*β*CD, have shown a specific affinity for sequestering 7KC and eliciting efflux of 7KC from cells, including macrophages and foam cells (Kritharides et al. (1996)). While CDs are generally used in pharmaceuticals as excipients for drug formulation, we propose that they could be used as an active pharmaceutical ingredient (API) themselves for the treatment of atherosclerosis and other dyslipidemias. CD derivatives such as HP*β*CD, SB*β*CD, and HP*γ*CD have been included in the FDA’s list of Inactive Pharmaceutical Ingredients due to their strong safety profile, and *β*CD has an optimal cavity diameter for encapsulating steroid molecules (such as cholesterols). We have engineered UDP-003, a CD dimer, to have not only the ideal cavity diameter, but also the correct cavity length to encapsulate sterols. We have further functionalized this molecule to create 300-fold specificity for 7KC over cholesterol (Anderson et al. (2024)).

**Figure 1:**
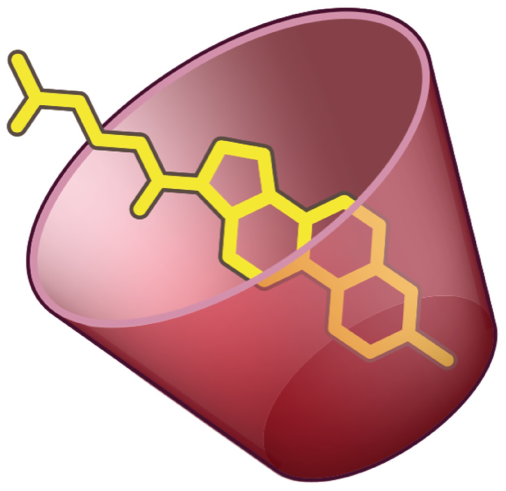
Host-guest inclusion complex with CD. Visual representation of the interaction of a guest with the hydrophobic cavity of a CD in a host-guest inclusion complex.

This paper aims to elucidate the mechanism by which UDP-003 is able to reverse foam cells back into healthy macrophages. While it has been demonstrated that administering high doses of HP*β*CD to hypercholesterolemic mice leads to decreased inflammation and plaque size (Zimmer et al. (2016)), UDP-003 has the potential to show much greater effects. Foam cell formation was induced in both mouse and human macrophages using 7KC or oxLDL, demonstrating that these treatments were sufficient to induce this morphological change. Furthermore, it was demonstrated that the application of UDP-003 reversed the negative effects on gene expression, phagocytic function, and ROS production. This data shows that UDP-003 and the specific targeting of 7KC mitigate and alleviate the foam cell phenotype, causing clearance of lipids, downregulation of ROS, and improvement of phagocytic activity. These findings suggest that UDP-003 is a promising potential therapeutic agent for treatment and prevention of atherosclerosis.

## 2. Materials and Methods

### 2.1. Materials

The following chemicals were used in the experiments: Dulbecco’s Modified Eagle’s Medium (DMEM) with high glucose was purchased from Corning. BioReagent penicillin-streptomycin 100X sterile mixture, DMSO, 2’,7’-Dichlorofluorescin diacetate (DCF-DA), Phorbol 12-myristate 13-acetate (PMA), 2-Mercaptoethanol, and Oil red O (ORO) powder were obtained from Sigma Aldrich. Fetal Bovine Serum (FBS) was purchased from Tissue Culture Biologicals. RAW 264.7 macrophages, THP-1 monocytes, and RPMI-1640 media were acquired from the American Type Culture Collection. 7-ketocholesterol (7KC) was purchased from Cayman Chemicals. Flat-bottomed hydrophilic 96 well plates were bought from BrandTech. OxLDL was obtained from Lee BioSolutions. RNA was extracted using the E.Z.N.A Total RNA Kit I with DNase I treatment, both of which were purchased from Omega Bio-Tek. Oil red O staining images were captured using ThermoFisher’s EVOS M5000 imaging system.

### 2.2. Cell Culture and Treatment

#### 2.2.1. Cell Culture

RAW 264.7 mouse macrophages were cultured in DMEM-high glucose (pH 7.2–7.4) supplemented with 10 % fetal bovine serum (FBS) and antibiotics (1 % penicillin-streptomycin). These cells were grown in a humidified atmosphere with 5 % CO2 at 37*^◦^*C. Cells were passaged every 2-3 days and were used for up to 15 passages. RAW 264.7 cells were plated at a density of 0.5*x*10^6^ cells/well in a 12-well plate. They were stimulated using either 7KC in ethanol or oxLDL in PBS for conditions requiring lipid loading. Cells were then incubated with 7KC or oxLDL for 24 to 48 h, depending on the modality.

THP-1 human monocytes were cultured in RPMI-1640 media supplemented with 10 % FBS, 1 % penicillin-streptomycin antibiotics mixture, and 100 *µ*M 2-mercaptoethanol in a humidified atmosphere with 5 % CO2 at 37*^◦^*C. The monocytes were passaged every 2-4 days and used until passage 25. Monocytes were seeded at a density of 1.5*x*10^6^ cells/ml in 6-well plates in growth media supplemented with 0.0185 *µ*g/ml PMA to initiate the differentiation process. Plates were incubated for 3 days to allow the monocytes to differentiate into macrophages. After incubation, the media was replaced with fresh media and the macrophages were further incubated for 24 h. The differentiated macrophages were stimulated with 7KC dissolved in ethanol for use in multiple assays.

#### 2.2.2. Treatment Modalities

Two modalities were used for 7KC and UDP-003 treatments: preventative and additive. In the preventative approach, 7KC and UDP-003 were added to the macrophages concomitantly for 48 h. In the additive approach, the cells were exposed to 7KC for an initial incubation of 24 h. Then, UDP-003 was introduced to the media containing 7KC and the solution was maintained for the remaining 24 h. 7KC was added at a concentration of 40 *µ*M for most assays or 10 *µ*M for the ORO assay. A lower 7KC concentration for the lipid analysis was used to minimize the lethal effects of 7KC while still inducing the morphology attributed to foam cells. 40 *µ*M 7KC was used for phagocytosis, ROS, and gene expression experiments, as this dose induces a severe cellular stress response, including apoptosis and necrosis, without being completely lethal to the cells. UDP-003 was administered at concentrations of 25, 50, 100, and 150 *µ*M.

### 2.3. Quantification of intracellular lipids

Intracellular oil accumulation was determined using the Oil red O (ORO) staining method. To prepare the ORO stock solution, ORO powder was dissolved in isopropyl alcohol in a ratio of 3:1 w/v. The solution was mixed on a rotator for 10 min, then incubated at room temperature for 10 min. After sedimentation of the particulates in the ORO stock solution, a working stock solution was subsequently prepared in a ratio of 3:2 v/v in deionized water. This mixture was further filtered down with a Steriflip filter to yield the ORO working stock.

The treated RAW 264.7 cells were washed with PBS and fixed with 4 % paraformaldehyde for 1 h. After the incubation, paraformaldehyde was aspirated and the cells were washed with 60 % isopropyl alcohol and air-dried before staining. Once dried, the macrophages were stained with ORO working solution for 5 min. The cells were then washed with water and images of the stained cells were taken using the EVOS M5000 imaging system. The ORO dye was eluted using 100 % isopropanol and the absorbance was measured with the SpectraMax i3 spectrophotometer at 510 nm.

### 2.4. Measurement of Phagocytosis Activity

The preventative and additive modalities were tested using these cells. The preventative modality consisted of the simultaneous incubation of 7KC and UDP-003 for a total of 48 h. The additive modality consisted of incubating cells with 7KC overnight, followed by the addition of UDP-003 for another 24 h. THP-1 cells were seeded in 6 well plates and were differentiated with PMA (0.0185 *µ*g/mL) followed by incubating the cells for 72 h. As previously described, additive modality was tested in THP-1 cells.

For both cell models, conditions were tested in triplicate and included positive and negative controls in the form of an equal volume of 7KC or ethanol, respectively. 7KC was used at a concentration of 40 *µ*M and UDP-003 treatment was used at concentrations of 100 *µ*M and 150 *µ*M for RAW 264.7 and 100 *µ*M for THP-1 macrophages.

Phagocytic activity was measured using the green *E.coli* assay kit from BioVision. Both macrophage cell lines were incubated with green *E.coli* diluted to 1 % in media for 2 h at 37*^◦^* C and the cells were harvested following manufacturer’s instructions. Each sample was stained with PI diluted to 1 % by buffer solution. The quantification of phagocytosis activity was determined by the amount of macrophages that internalized fluorescently pre-labeled *E.coli* particles, using the BD Accuri C6 flow cytometer.

### 2.5. Preparation of total RNA and gene expression analysis (Real-Time RT-PCR)

Total RNA was extracted after treatments using an Omega Biotek RNA Extraction Kit with DNase I treatment. RNA was quantified using a NanoDrop ND-2000 spectrophotometer. 0.5 *µ*g of RNA was reverse transcribed into cDNA using the Primescript*^TM^* RT Master Mix and following the manufacturer’s protocol. The cDNA, combined with Luna Universal qPCR Master Mix and 5pmol of the primer of interest, was amplified using Roche LightCycler 96 qPCR machine. For internal control, the GAPDH gene was used for normalization of each gene of interest and the relative expression was determined through the Ct method (ΔΔCt), also known as the 2-ΔΔCt method. Sequences of the primer used are listed in Supplementary Table 1.

### 2.6. Detection of intracellular ROS production

Intracellular ROS production was evaluated using 2,7’-Dichlorofluorescin diacetate (DCF-DA). The cell culture media was aspirated and cells were exposed to 10 *µ*M DCF-DA in PBS for 25 min. After incubation, the supernatant was aspirated and the cells were washed with PBS. The cells were harvested and centrifuged. The pellet obtained after centrifugation was washed with PBS thrice. Cells were resuspended in PBS, and PI was added to sort out viable cells. Fluorescence was measured using flow cytometry (BD Accuri flow cytometer) with an excitation wavelength of 485 nm and an emission wavelength of 535 nm. The relative fold changes were calculated compared to the negative control and intracellular levels of ROS were determined by taking the raw cell count and dividing it by the average cell count for the untreated (EtOH) condition.

### 2.7. Detection of Apoptotic and Necrotic cells

For this set of experiments, the THP-1 macrophages were treated with 7KC and UDP-003 simultaneously for 48 h. Measurement of apoptosis and necrosis was carried out following the instructions from the Apoptosis/Necrosis Assay Kit from Abcam. Supernatant from the treated THP-1 macrophages was collected. The adhered macrophages were washed with sterile PBS and the cells were harvested and collected along with the supernatant. The tubes were centrifuged and the supernatant was discarded. The pellet was washed with PBS, centrifuged for 5min, and then resuspended in the assay buffer. The cells were further stained with Apopxin Green Indicator for apoptosis measurements and 7-AAD for necrosis measurements. The macrophages were incubated at room temperature for 30 min. The samples were diluted with assay buffer after incubation, as per instructions in the manual, and analyzed in the BD Accuri flow cytometer. For apoptosis quantification, the stained cells were detected with an excitation wavelength of 533 nm and an emission wavelength range of 503-563 nm (FL-1 filter), while necrosis readings were at an excitation wavelength of 670 nm (FL-3 filter). Measurements for apoptotic cells were quantified using mean FL-1H fluorescence.

### 2.8. 7KC extraction from plaque-containing human tissue using UDP-003

Human vascular tissue samples containing plaque were sliced into pieces of 5 to 50 mg, and were transferred into individual 2 ml Eppendorf tubes. Samples were treated with HPBCD monomer in PBS (0.5 mM, 1 mM, 2.5 mM, or 5 mM), or UDP-003 in PBS (0.25 mM, 0.5 mM, 1 mM, or 2.5 mM) with equivalent PBS control in a ratio of 1:10 w/v. Samples were shaken in the incubator at 37*^◦^*C for either 15 min, 3 h, or 24 h. Subsequently, the supernatant from the human tissue was transferred to separate tubes for liquid-liquid sterol extraction.

Human plaque tissue required an extra homogenization step before liquid-liquid extraction. The tissue was homogenized in the presence of stainless steel beads, which facilitate homogenization, along with sterile water. Beads and sterile water were added in the ratio of 9:1 v/w tissue. The tissue samples were homogenized in a Next Advanced Bullet Blender for 5 min and transferred into clean Eppendorf tubes. To the supernatant collected previously and the homogenized human tissue, an equal volume of ethanol as the aqueous solution, 200 *µ*g/ml of butylated hydroxytoluene (BHT), and hexane, at a 2:1 ratio relative to the amount of ethanol, were added. The sample tubes were placed in a tube shaker at 1500 rpm for 1 h. After shaking, the samples were centrifuged for 5 min at room temperature and 90 % of the hexane layer was transferred into glass vials. Fresh hexane was added back into the glass vials and the extraction was repeated, with samples shaken on the tube shaker for an additional 5 min, followed by centrifugation for 5 min. The hexane layer containing the oxysterols was aliquoted and transferred into a glass vial. Samples were dried with a gentle stream of nitrogen gas and stored at -80*^◦^*C until 7KC quantification using LCMS.

### 2.9. Measurement of viability

Cells were plated at a density of 0.5*x*10^6^ cells/well in a 12-well plate and allowed to adhere overnight in the presence of complete growth media. The cells were then treated with different concentrations of UDP-003 (10, 100, 200, 300 *µ*M). The negative control was treated with an equivalent volume of PBS used to dissolve the UDP-003 treatment. The cytotoxic effects were measured using propidium iodide (PI). PI penetrates and disrupts the plasma membrane and stains the nucleic acids of nonviable cells (Zhao et al. (2010)). Cells were harvested after 24 h of treatment and were pelleted down. Subsequently, the cells were washed two successive times with PBS. Finally, the pellet was reconstituted in PBS and 1 % PI. The mixture was then measured using a BD Accuri C6 flow cytometer with excitation maxima of 535 nm and an emission maxima of 615 nm.

### 2.10. Statistical Analysis

All datasets are expressed as the mean of at least three independent replicates and results were compared by one-way analysis of variance (ANOVA) using Prism 9.3.0.

## 3. Results

### 3.1. UDP-003 prevents and reverses the accumulation of intracellular lipids in RAW 264.7 cells

Lipid droplet accumulation is the most distinctive morphological feature of foam cells. Cells were stained with ORO to determine and compare the relative amount of lipid droplets in the 7KC-induced foam cells before and after UDP-003 treatment, and the staining was measured by UV-Vis spectrophotometry. When foam cells were concurrently treated with UDP-003 and 10 *µ*M 7KC, a reduction in absorbance of 13.3 % was observed relative to the absorbance seen with the foam cells that were treated with 7KC (Fig. 2A,C). This reduction in absorbance can be explained by the ability of UDP-003 to sequester the 7KC before it can enter the cell and cause cellular dysfunction. A reduction in absorbance was also observed when the foam cells were treated with UDP-003 (in the presence of 7KC) for 24 h after already being exposed to 7KC for 24 h (Fig. 2B,D). In this modality, the absorbance dropped by 29.2 % when treated with 100 *µ*M UDP-003. These data, combined with our previous findings that our proprietary CDs can remove 7KC from foam cells, indicates that UDP-003 can not only prevent the foam cell-inducing effects of 7KC, but can reverse foam cell formation once it has already occurred.

**Figure 2:**
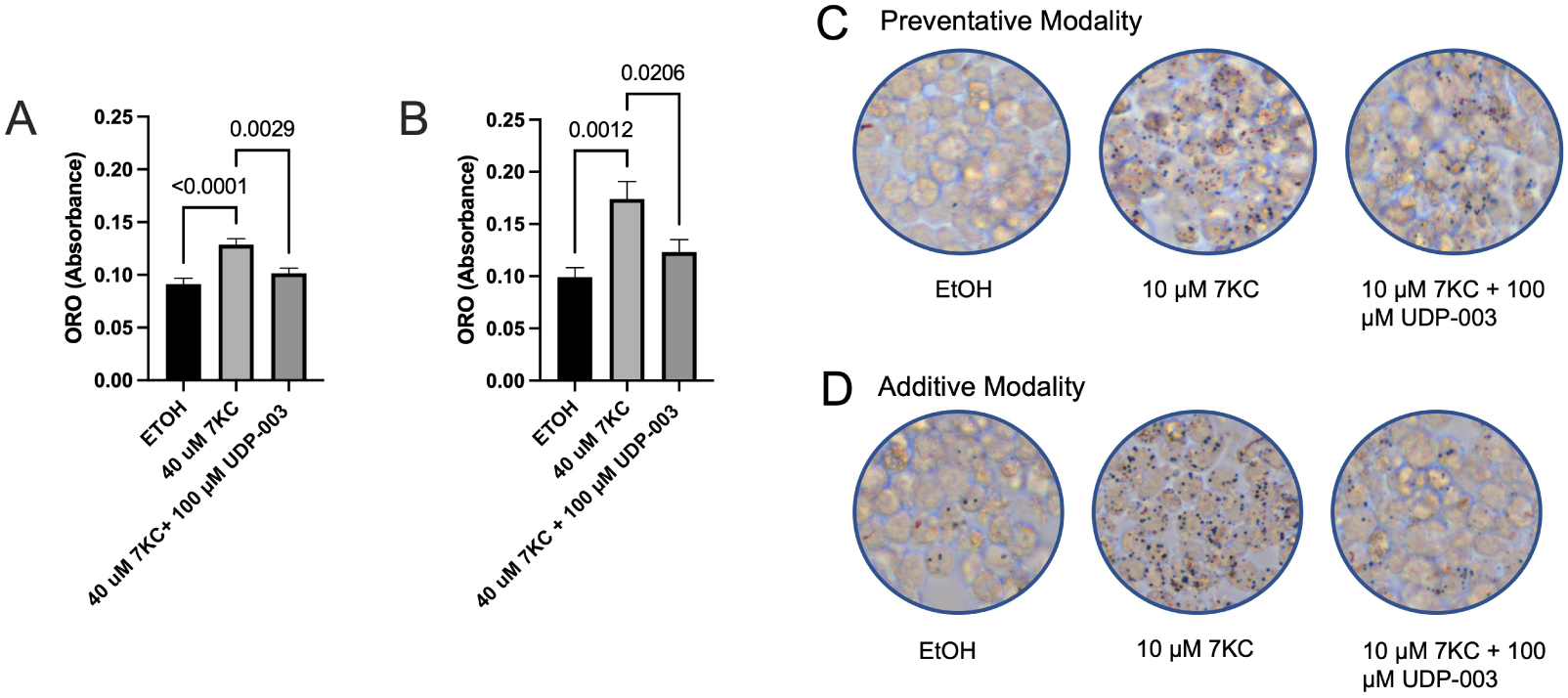
In vitro reduction of intracellular lipid droplets in 7KC-treated RAW 264.7 macrophages with UDP-003. Quantification of intracellular lipid droplets collected from RAW 264.7 macrophages was through Oil Red O. Under all modalities, the positive control macrophages were treated with 10 *µ*M 7KC for 48 h. The macrophages were treated under (A) preventative modality and (B) additive modality. Data is expressed in average absorbance with the baseline absorbance subtracted *±* SE bars from at least three independent replicates. Statistical analysis was performed using a one-way analysis of variance (ANOVA). Photos of the RAW 264.7 macrophages stained with Oil Red O under (C) pre-ventative modality and (D) additive modality were taken with the EVOS microscope.

The intracellular lipid droplet formation in foam cells that were induced by oxLDL was also evaluated as a potential physiological source of 7KC. As seen in Supplementary Fig. 1A, foam cells that were concurrently treated with 150 *µ*g/ml oxLDL and 200 *µ*M UDP-003 had a 31.4 % decrease in absorbance relative to foam cells only treated with 150 *µ*g/ml oxLDL. For foam cells that were exposed to 150 *µ*g/ml oxLDL for 24 h and then treated with 100 *µ*M UDP-003 in new media, Supplementary Fig. 1B shows that when stained with Oil Red O, the treated cells show 17 % less absorbance than that seen in foam cells that were exposed only to oxLDL for 48 h. Supplementary Fig. 1C shows a decrease in absorbance of foam cells that were treated with UDP-003 for 24 h while being exposed to oxLDL for 48 h, relative to the absorbance from cells that were only treated with oxLDL during the 48 h period. Therefore, UDP-003 can prevent and reverse the foam cell morphology induced by both 7KC and oxLDL.

### 3.2. UDP-003 increases the phagocytosis activity of THP-1 and RAW 264.7-derived foam cells

In addition to being laden with lipid droplets, foam cells are characterized by their “quasi-senescence.” This includes an inability to engulf noxious entities within the body, the primary function of macrophages. To determine the efficacy of UDP-003 in preventing and rescuing loss of phagocytic function in macrophages due to exposure to 7KC, THP-1 and RAW 264.7 macrophages were treated under preventative and additive modalities as described in the material and methods section. Treatment with UDP-003 in the preventative modality prevented the loss of phagocytic activity when cells were exposed to 7KC (Fig. 3B). Simultaneous incubation of 7KC and UDP-003 in RAW 264.7 macrophages, compared to the 40 *µ*M 7KC control, resulted in 14.5 % increased uptake of green E.coli particles for the 100 *µ*M UDP-003 preventative treatment. A 150 *µ*M UDP-003 treatment resulted in a 18.9 % increase in phagocytic ability (Fig. 3B). This indicates recovery of phagocytic ability. In the additive modality, UDP-003 rescued the phagocytic ability of RAW 264.7 by 10.8 % at 100 *µ*M and 18.2 % at 150 *µ*M (Fig. 3C). Experiments in RAW 264.7 cells show a dose dependent recovery of phagocytic ability with an increase in UDP-003 concentration (Fig. 3A). Similarly, THP-1 macrophages showed a 12.1 % increase in phagocytic ability when treated with 100 *µ*M UDP-003 under the additive modality (Fig. 3D). Thus, UDP-003 is able to reverse the deleterious effects of 7KC of the phagocytic function of foam cells derived from THP-1 and RAW 264.7 cells.

**Figure 3:**
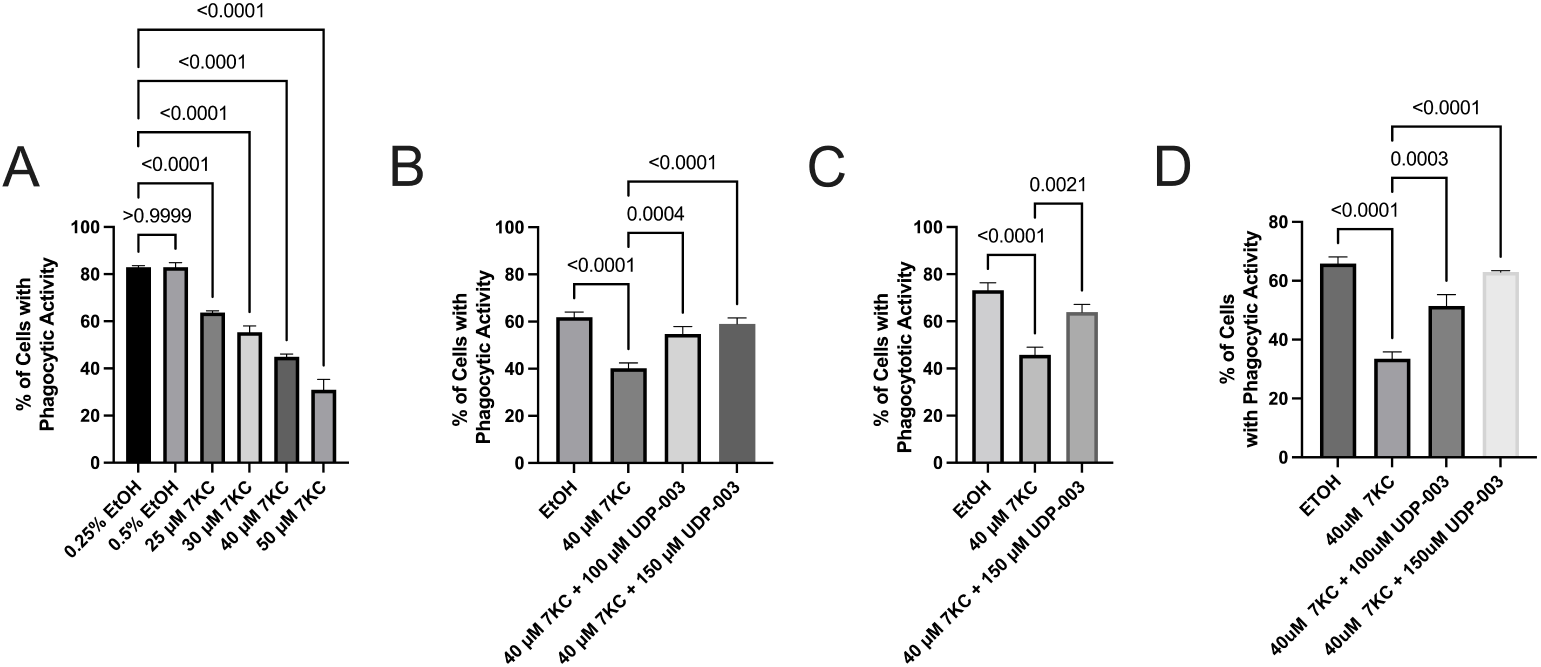
In vitro restoration of phagocytic activity with UDP-003 against 7KC treated macrophages and effects of various 7KC concentration to phagocytic ability. Phagocytosis activity of both RAW 264.7 and THP-1 macrophages was measured through the intake of green fluorescently prelabeled E.coli particles in the BD Accuri flow cytometer. (A) RAW 264.7 macrophages were exposed to various concentrations of 7KC (25-50 *µ*M) with the equivalent volume of EtOH as negative control to determine concentrations that can be used as positive controls. Under all modalities, the positive control macrophages were exposed to 40 *µ*M 7KC for 48 h. The macrophages were treated under (B) the preventative modality and under (C) the additive modality. (D) The THP-1 macrophages were treated under the additive modality. Data is expressed in the mean percentage of cells that displayed phagocytic activity through the engulfment of the green E.coli particles *±* SE bars. The data was collected from at least three independent replicates. Statistical analysis was performed using a one-way analysis of variance (ANOVA).

### 3.3. UDP-003 downregulates the expression of PPARγ and MIP1α genes in RAW 264.7 cells

To elucidate the underlying mechanism by which UDP-003 rescues the morphological and functional phenotype of foam cells, mRNA levels for peroxisome proliferation-activated receptor gamma (PPAR*γ*), Lectin-like oxidized low-density cholesterol lipoprotein receptor 1 (LOX-1), and Macrophage Inflammatory Protein 1 alpha (MIP1*α*) were measured by qRT-PCR. The experimental setup involved exposing RAW 264.7 cells to 40 *µ*M 7KC for 48 h. A substantial upregulation of all three genes was observed with treatment with 40 *µ*M 7KC (Fig. 4). PPAR*γ* exhibited a 1.8-fold increase, while MIP1*α* increased 5-fold, and LOX-1 increased 22-fold. Treatment with UDP-003 at concentrations of 25 or 100 *µ*M downregulated the expression of MIP1*α* (2-fold) and PPAR*γ* in the additive modality wherein UDP-003 was added to the cells after 24 h of treatment with 7KC. When cells were treated concomitantly with UDP-003 and 7KC, a dose-dependent decrease was observed for LOX-1; however, this decrease was not statistically significant. Moreover, no significant differences were observed when UDP-003 was added after 24 h treatment with 40 *µ*M 7KC. For MIP1*α*, a decrease of 2.6-fold and 1.3-fold were observed at 25 *µ*M and 100 *µ*M UDP-003 respectively. Combined, these data indicate that UDP-003 is able to reverse some aspects of the shift in gene expression induced by 7KC.

**Figure 4:**
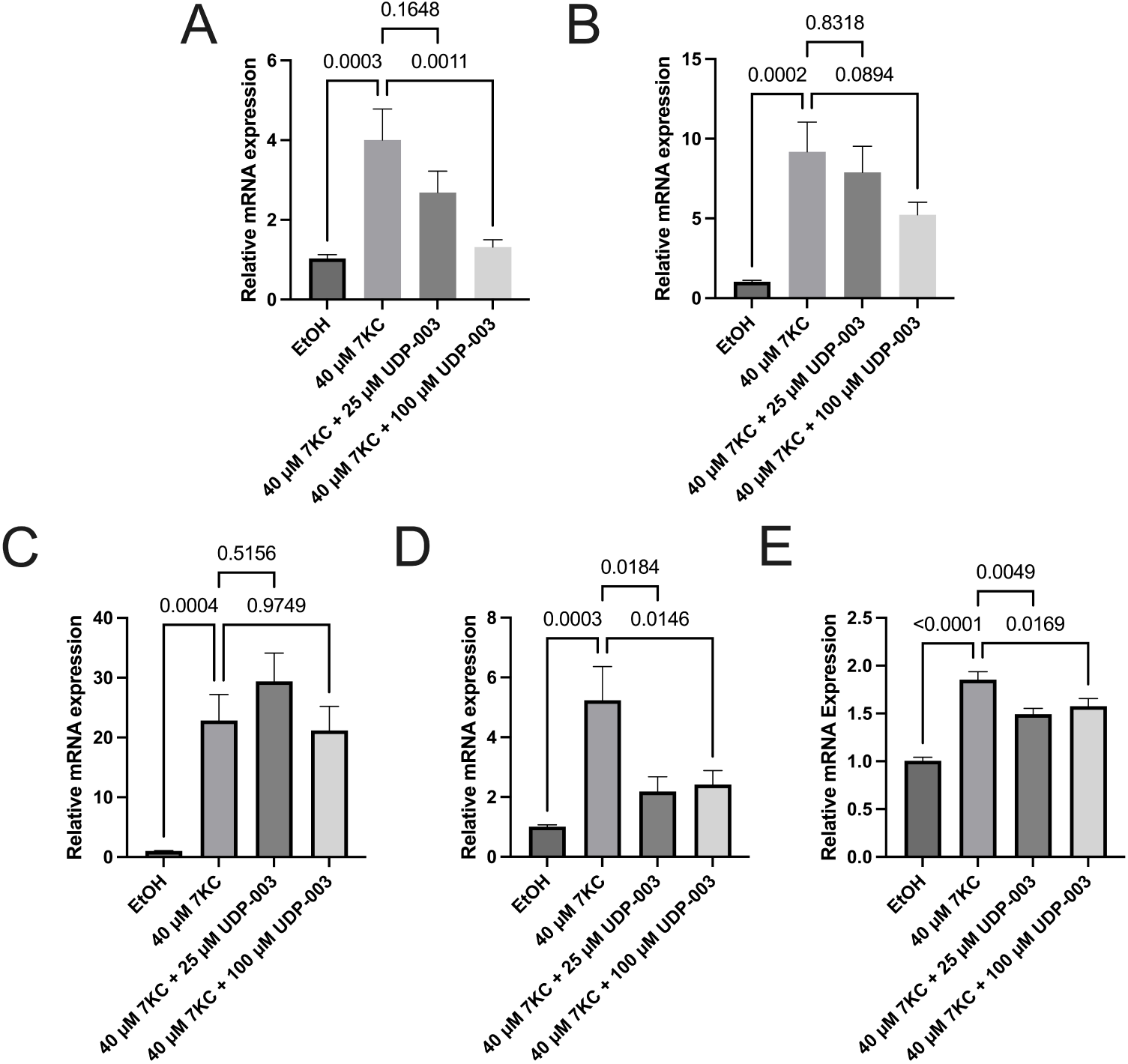
In vitro reduction of PPAR*γ*, MIP1-*α* gene expression by UDP-003 against 7KC treated RAW 264.7 cells. qPCR analysis of mRNA expression levels represented relative to the control of the experiment, RAW 264.7 cells were seeded at a density of 0.5 x 106 cells/well in a 12-well plate and were stimulated by 40 *µ*M 7KC for 48 h in preventative modality (A, B), and in additive modality (C, D, E). Subsequently, total RNAs were prepared and subjected to qRT-PCR analysis using specific primer sets (Supplementary Table 1), responsible for MIP1*α* (A, D), LOX-1 (C) and PPAR*γ* (B, E). The genes were normalized to GAPDH. The negative control is treated with equivalent EtOH and PBS. Data obtained from at least triplicate repeats are shown as mean *±* SE. Statistical analysis was performed using a one-way analysis of variance (ANOVA).

### 3.4. UDP-003 decreases inflammation by reducing the levels of intracellular ROS in RAW 264.7 cells

To further demonstrate the efficacy of UDP-003 in reducing the inflammatory effects of 7KC, intracellular ROS levels were measured in RAW 264.7 macrophages under the preventative and additive modality as described in the Materials and Methods section. In the preventative modality, 40 *µ*M 7KC led to a 3.53-fold increase in intracellular ROS induced fluorescence relative to untreated macrophages. Compared to macrophages treated with 40 *µ*M 7KC, macrophages treated with 7KC concurrently with 25 *µ*M UDP-003 had a 1.32-fold reduction in ROS-induced fluorescence. Notably, the addition of 100 *µ*M UDP-003 induced a 2.27-fold decrease in ROS levels (Fig. 5A). Similarly, in the additive modality, 7KC-treated macrophages exhibited a 5.79-fold increase in ROS induced fluorescence compared to untreated macrophages (Fig. 5B). Upon the addition of 25 *µ*M and 100 *µ*M UDP-003 24 hours after the 7KC treatment, macrophages exhibited a 1.27 fold and 2.0-fold decrease in ROS levels, respectively (Fig. 5B). This indicates that UDP-003 effectively ameliorated intracellular ROS concentrations in macrophages deleteriously impacted by 7KC in a dose-dependent manner.

**Figure 5:**
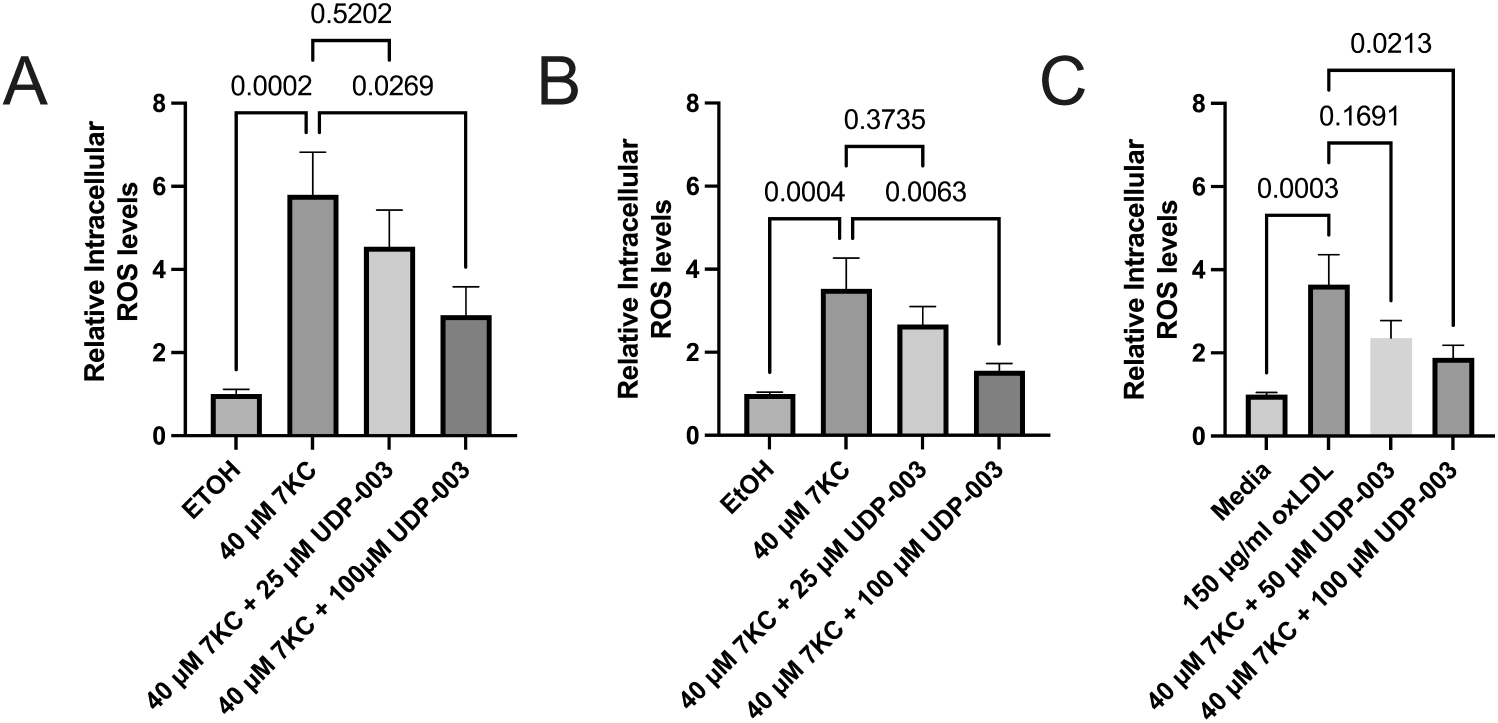
In vitro reduction of intracellular ROS production with UDP-003 against 7KC treated macrophages. The measurement of intracellular ROS levels demonstrated a significant increase in RAW 264.7 macrophages treated with 40 *µ*M 7KC in the preventative modality (A) and in the additive modality (B). Macrophages were treated with both 7KC and 25 *µ*M UDP-003 as well as 100 *µ*M UDP-003. Data are expressed as relative fold changes. Data obtained from at least triplicate repeats are shown as mean *±* SE. Statistical analysis was performed using a one-way analysis of variance (ANOVA).

Relative to untreated cells, oxLDL-treated macrophages exhibited a 3.64-fold increase in ROS levels (Fig. 5C). Compared to oxLDL treated cells, macrophages treated with both oxLDL and 50 *µ*M UDP-003 demonstrated a 1.54-fold decrease, while 100 *µ*M UDP-003 treated macrophages exhibited nearly a 1.93-fold decrease in ROS levels (Fig. 5C). The addition of UDP-003 to macrophages had a similar effect on intracellular ROS levels induced by oxLDL to macrophages treated with 7KC, with a significant decrease at 100 *µ*M UDP-003.

### 3.5. UDP-003 mitigates apoptosis and necrosis in THP-1 cells

Over the course of atherogenesis, macrophages become foam cells which then undergo apoptosis and, eventually, necrosis. Late stage atherosclerotic lesions are characterized by a substantial necrotic core composed mostly of dead foam cells. To explore the effects of UDP-003 on this process, THP-1 macrophages were treated with 100 *µ*M UDP-003 and 40 *µ*M 7KC for 48 h and a decrease of 39.6 % in mean FL-1H fluorescence was observed (indicative of apoptotic cells), relative to the macrophages that were only treated with 7KC (Fig. 6A). Meanwhile, upon the simultaneous treatment of UDP-003 and 7KC, the amount of necrotic cells decreased by 58.7 % when compared to the macrophages treated with only 7KC (Fig. 6B). Based on these results, UDP-003 can prevent foam cells from undergoing apoptosis and necrosis, indicating the potential to prevent the progression of atherosclerotic plaques to the stages at risk of rupture and thrombosis.

**Figure 6:**
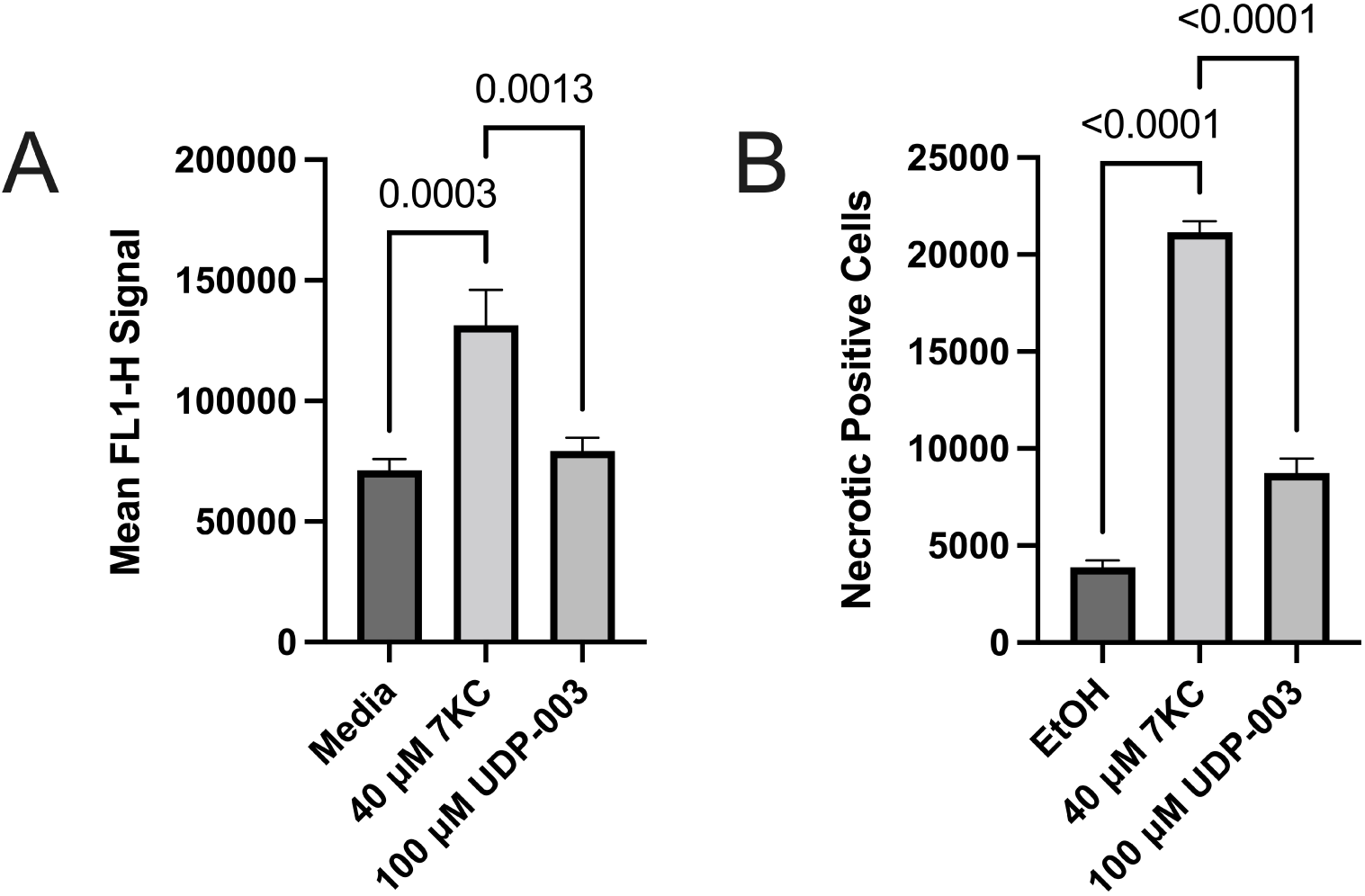
In vitro prevention of apoptosis and necrosis in 7KC-treated THP-1 macrophages with UDP-003. Quantification of mean fluorescence from the FL-1 channel and percentage of necrotic positive cells from THP-1 when the cells are stained with Apopoxin Green for apoptosis (A) and 7-AAD for necrosis (B). The macrophages were stained with the aforementioned reagent for 30min before reading in the BD Accuri flow cytometer. Data for apoptosis is expressed in average FL-1H fluorescence *±* SE bars from at least three independent replicates. Data for necrosis is expressed in the average percentage of THP-1 macrophages that are considered “necrotic” based on a gating for 7-AAD staining *±* SE bars from at least three independent replicates. Statistical analysis was performed using a one-way analysis of variance (ANOVA).

### 3.6. UDP-003 exhibits no impact on the viability of RAW 264.7 cells

As with any potential therapeutic, it is important to demonstrate that UDP-003 is not toxic to the targeted cells. UDP-003 exhibited no significant cytotoxic activity at concentrations of up to 300 *µ*M, as shown in Supplementary Fig. 2 It is important to note that at 300 *µ*M, UDP-003 displayed slight toxicity, with cell viability dropping by 19 %. Based on these results, we avoided further use of 300 *µ*M. As depicted in the figure, when cells were exposed to up to 200 *µ*M UDP-003, their levels of cell viability were similar to those of the non-treated control group. For this study, the cell viability results were expressed as a percentage of normalized non-treated cells, and the experiments were executed in triplicate.

### 3.7. UDP-003 successfully removed 7KC from human vascular tissue

We have previously shown the ability of our proprietary CDs to remove 7KC from foam cells in vitro. The foam cells involved in atherosclerosis exist, however, in plaques within the arterial wall. In order to demonstrate the ability of UDP-003 to remove 7KC from the tissue in which it causes pathology, we attempted to pull 7KC out of ex vivo human plaque-laden arterial tissue. Human arterial tissue, confirmed to contain atherosclerotic plaque, was incubated with the HP*β*CD monomer or UDP-003 dimer at various concentrations for 15 min, 3 h, and 24 h. Because UDP-003 is dimerized, the treatment was administered at half the concentration range of the HP*β*CD monomer for the 15 min and 3 h conditions. Across many of the incubation periods, UDP-003 treatment showed a greater percentage of free 7KC removal compared to the corresponding HP*β*CD monomer treatment. Incubation in PBS alone (mock) for 15 min showed a baseline reduction of 4.03 % from the untreated plaque tissue. The 15 min treatment with 5 mM HP*β*CD removed 21.06 % of 7KC, while 2.5 mM UDP-003 treatment removed 27.30 % of 7KC (Fig. 7). When treated for 3 h, 5 mM HP*β*CD extracted 15.84 % of 7KC, while 1 mM UDP-003 extracted 17.27 % and 2.5 mM UDP-003 removed 31.34%. After 24 h of treatment, administration of 2.5 mM HP*β*CD removed 6.8% and 2.5 mM UDP-003 removed 27.05 %. Treatment with 2.5 mM HP*β*CD showed that a lower incubation period elicited a smaller percentage of 7KC removed. These results suggest that the dimerized CD has increased binding affinity for 7KC compared to monomeric CD, and has increased ability to solubilize 7KC. This has been attributed to the formation of non-covalently-bonded 1:1 inclusion complexes in the dimerized CD (Anderson et al. (2021, 2024)). Additionally, these experiments demonstrate that the time-constant for 7KC removal from intact plaque-laden arterial tissue is at or below 15 min, indicating that systemic administration of UDP-003 is likely to allow ample exposure of plaques to UDP-003 to achieve meaningful 7KC removal.

**Figure 7:**
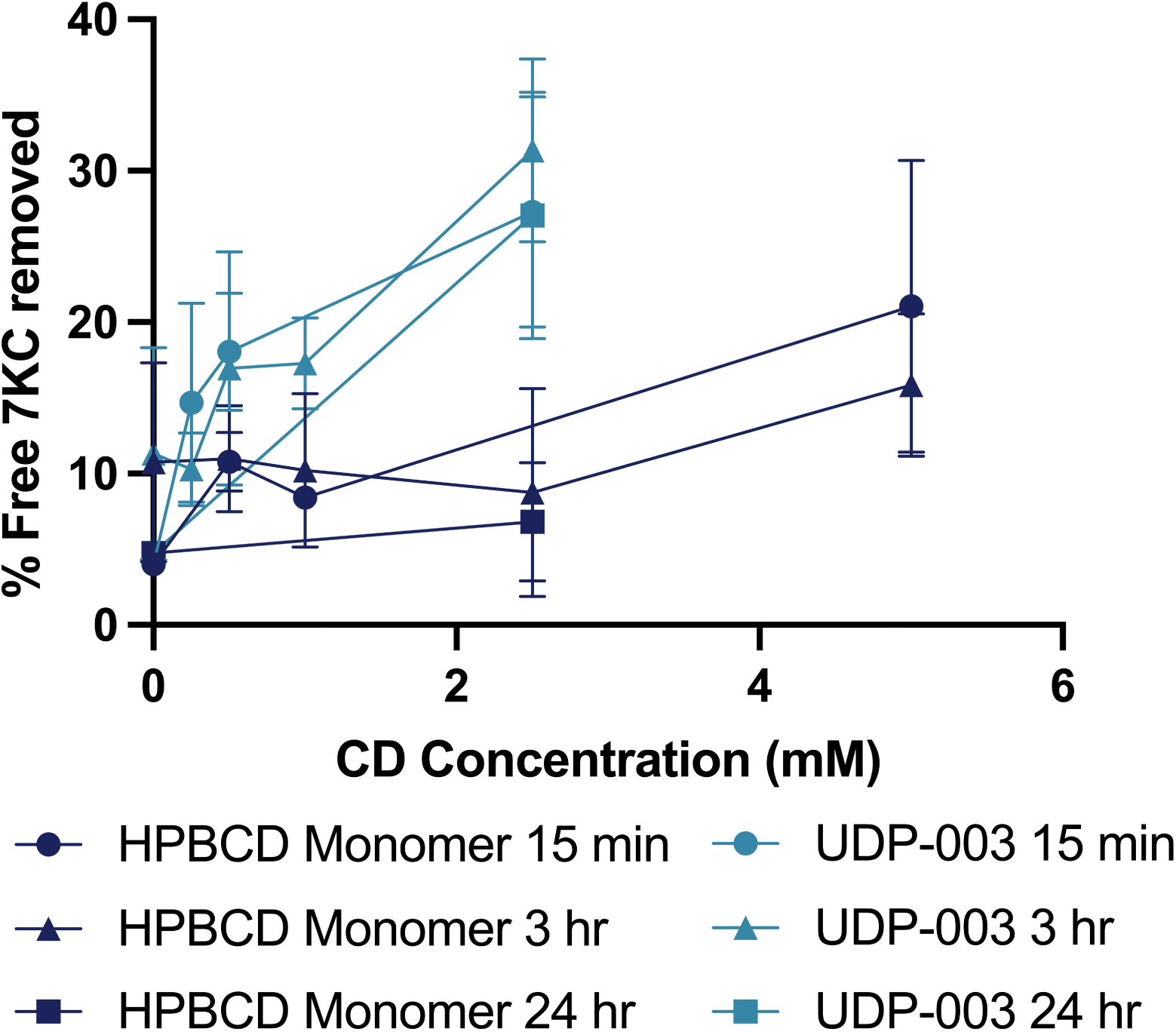
Ex vivo 7KC extraction from plaque-containing human tissue. Quantification of 7KC removed from plaque-containing human tissue samples after exposure to aqueous solutions with either HP*β*CD (dark blue lines) or UDP-003 (light blue lines). The human tissue samples were exposed to the CD solutions for either 15 min (circle markers), 1 h (triangle markers), and 3h (square markers). Data is expressed as the percentage of 7KC extracted into the treatment solution, relative to the total amount of 7KC found in the tissue and solution. *±* SE bars.

## 4. Discussion

Lipid metabolism is a critical process that drives a variety of cellular functions (Florance and Ramasubbu (2022); Maxfield et al. (2023); Yan and Horng (2020)). While cholesterol is required for all animal cells, oxidation at the 7 position unleashes 7KC, a slow-killing poison that causes chronic inflammation, beginning with dysfunction in endothelial cells followed by excessive infiltration of monocytes at the site of inflammation. Subsequent increases in proinflammatory cytokines furthers the recruitment of cells to the site of inflammation, giving rise to a positive feedback loop. 7KC accounts for 30 % of the total cholesterol in oxLDL, and is an ever-growing presence in the body with no means of clearance (Brown et al. (1996)). In this study, well-established cell models have been used to test whether (1) treatment with 7KC is sufficient to induce foam cell formation and (2) whether removal of 7KC with UDP-003 can restore function to these macrophages. This could have implications for the treatment of atherosclerosis and other inflammatory diseases.

This study also highlights the use of different approaches to treat foam cells with UDP-003 to facilitate macrophage restoration. In the preventative method, it was hypothesized that UDP-003 prevents 7KC from entering the system by encapsulating it upfront, and minimizes 7KC contact with the cells. In the additive method, a real-world system was mirrored, as in a real-world situation, foam cell formation and plaque buildup is generally not discovered until after dysfunction is prevalent. Administration of UDP-003 after prolonged 7KC exposure simulates administration of treatment after disease progression.

Foam cell formation was successfully induced in both RAW 264.7 and THP-1 macrophages, well established cell models for atherosclerosis, by 7KC or 7KC-containing oxLDL over a period of 48 h. This treatment resulted in an increase in cytokine release, lipid accumulation, oxidative stresses, and impaired phagocytosis. Administration of UDP-003 mitigated these foam cell characteristics by encapsulating 7KC to reduce lipid accumulation and ROS, and improving the phagocytosis capability of the cells.

A plethora of studies have demonstrated that 7KC toxicity reduces oxidative phosphorylation, increases oxidative stress, and increases lipid accumulation. These effects further decrease the phagocytic ability of the cells, causing excessive ER stress and downregulation of phosphatidylinositol 4,5-bisphosphate (Calle et al. (2019); Játiva et al. (2022); Nury et al. (2018)). Our data show a decrease in phagocytosis after exposure to 7KC in THP-1 and RAW 264.7 cells. UDP-003 successfully improves phagocytosis in THP-1-derived foam cells and in RAW 264.7-derived foam cells in a dose-dependent manner. A dose-dependent decrease in phagocytosis with increasing concentrations of 7KC was also seen (Fig. 3A).

While UDP-003 dramatically restores phagocytosis in lipid-laden foam cells, it does not completely restore their activity to baseline levels. This could be explained by the fact that a high concentration of 7KC (40 *µ*M) drives the cells towards apoptosis and necrosis. As shown in THP-1 cells, exposure to 40 *µ*M 7KC for 48 h exerts a two-fold increase in apoptosis and a sixfold increase in necrosis. 7KC induces a cascade of events to disrupt homeostasis by these two characteristic modes of cell death. 7KC induces LC3II/LC3I ratio and SQSTM1/p62 expression in RAW 264.7 cells leading to autophagy (Wang et al. (2022)). Inhibition or reversal of the apoptotic and necrotic core formation would be one mechanism of reducing the progression of the necrotic core. UDP-003 ostensibly reverses and prevents further apoptosis by decreasing it by 40 % as compared to the 7KC treated control and decreasing the necrosis process by 60 % in THP-1 cells. Thus, UDP-003 is able to prevent apoptosis and necrosis from progressing.

This data also illustrates an evident elevation in intracellular ROS levels in 7KC induced foam cells, and presents evidence that administration of UDP-003 significantly reduces cellular levels of ROS. This decrease in ROS levels with UDP-003 treatment could reduce the amount of cholesterol oxidation into 7KC (or a precursor to 7KC) through ROS, closing the positive feedback loop of inflammation that 7KC causes (Fig. 8A).

**Figure 8:**
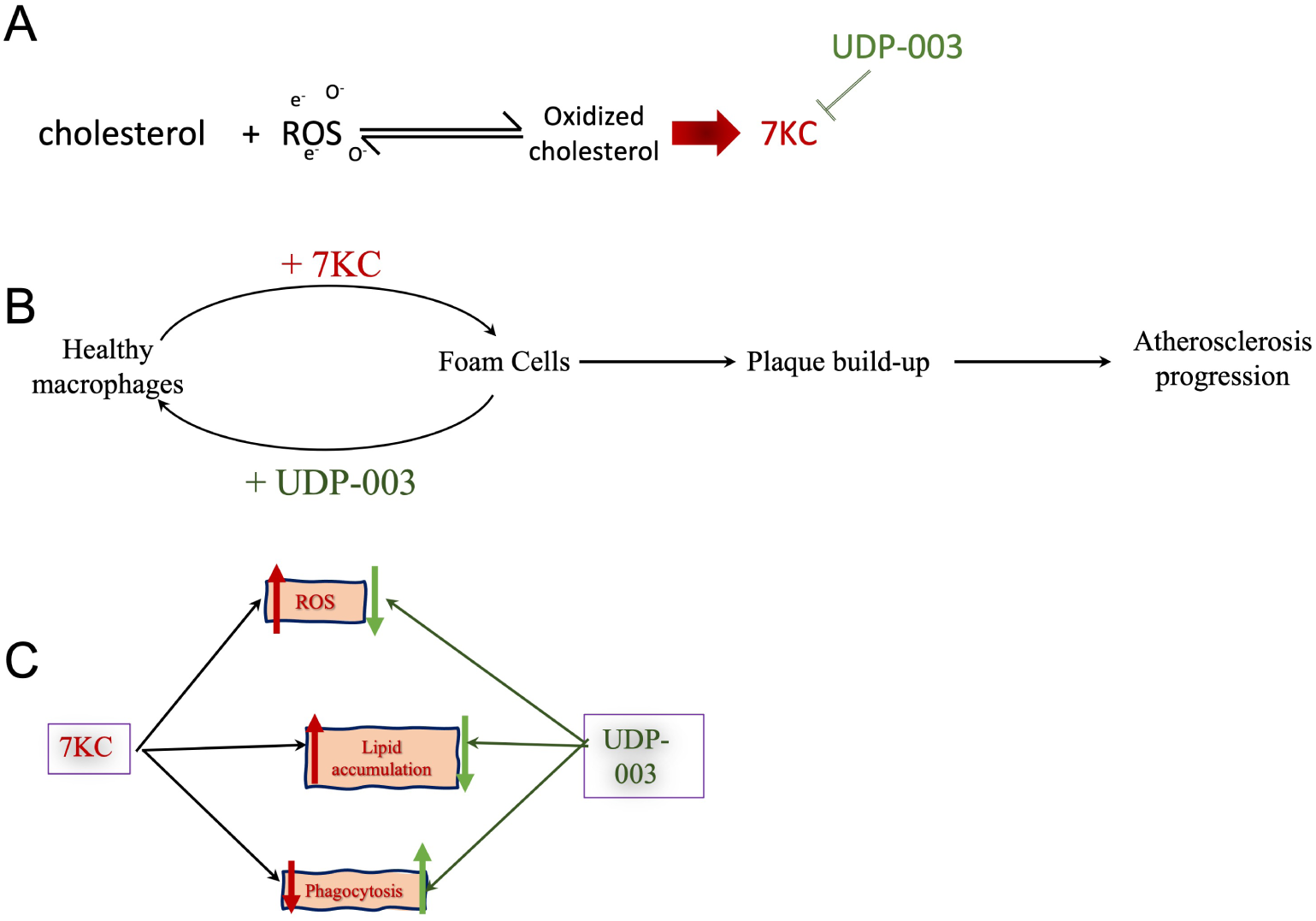
General mechanism model of the interactions of 7KC and UDP-003 with foam cells/macrophages. (A) Model of the oxidation of cholesterol with the presence of ROS into oxidized cholesterol including 7KC and the precursors to 7KC. UDP-003 sequesters 7KC, inhibiting the positive feedback loop to produce more 7KC. (B) Model of the atherosclerosis progression pathway of macrophages with 7KC converting healthy macrophages to foam cells and UDP-003 reverting the foam cells back into healthy macrophages. (C) Model of the effects of 7KC (red arrow) and UDP-003 (green arrow) to key characteristics of macrophages/foam cells.

Another hallmark of foam cell formation is the accumulation of lipids within macrophages. Macrophages ingest oxLDL, LDL, and 7KC by the processes of macropinocytosis, phagocytosis, or via receptors such as LDLR, LOX-1, and CD36. These lipids are metabolized in the lysosome, releasing free cholesterol and free fatty acids, which are further esterified in the endoplasmic reticulum into cholesteryl esters. Our data agrees with previously published work showing that 7KC exacerbates accumulation of intracellular lipid accumulation. UDP-003 significantly reduces lipid accumulation and effectively restores lipid levels to the baseline levels in RAW 264.7 cells.

The data shows upregulation of the chemoattractant gene MIP1*α* in the presence of 7KC. MIP1*α*, like MCP1, assists in recruiting immune cells (Bhavsar et al. (2015)). As 7KC is an important component of oxLDL, the impact of UDP-003 in downregulating LOX-1, a well established receptor for oxLDL, was investigated. Upregulation of LOX-1 was observed with addition of 40 *µ*M 7KC, however, UDP-003 did not suppress this upregulation.

Accumulation of cellular cholesterol activates LXR*α*, PPAR*γ*, SREBPs, NFkB, and multiple additional transcription factors (Neve et al. (2000); Remmerie and Scott (2018); Yu et al. (2015)). To further delve into the mechanism by which UDP-003 acts against 7KC, the mRNA expression of PPAR*γ* was measured. An upregulation of PPAR*γ* in the presence of 7KC was observed, and UDP-003 administration suppressed this expression at a dose of 100 *µ*M in the additive modality. This presents evidence that sequestration of 7KC by UDP-003 could reduce the 7KC-induced upregulation of PPAR*γ*.

In summary, we have designed and synthesized CD dimers and have shown that they have a much higher affinity to 7KC compared to monomeric CDs (Anderson et al. (2021, 2024)). In vitro experiments involving addition of 7KC to THP-1 human monocytes revealed that the screened dimers had the ability to sequester 7KC from the monocytes, minimizing 7KC’s cytotoxic effects towards these cells. In addition, many of the screened dimers show low hemolytic activity when added to whole blood (Anderson et al. (2021)). From the different CD dimers screened, one CD dimer (UDP-003) was designated as our lead candidate for further in vitro and in vivo experiments to test the candidate’s potential as a therapeutic drug. UDP-003 was able to restore phagocytic function to foam cells, reduce the upregulation of PPAR*γ*, prevent apoptosis and necrosis from further progressing, and successfully remove high concentrations of free 7KC from atherosclerotic plaque tissue.

## 5. Conclusion

This work presents several compelling conclusions regarding the mechanism of atherosclerosis progression as well as potential interventions. It is evident that exposure to 7KC or oxLDL can turn healthy macrophages into lipid-laden and dysfunctional foam cells, highlighting their significant role in atherosclerosis and disease progression. Impressively, treatment with UDP- 003 rejuvenates these foam cells, reviving their ability to phagocytose and fulfill their role of clearing and preventing plaque buildup. This suggests that UDP-003 has the potential to treat atherosclerosis via an entirely new mechanism. While more research is needed to fully understand these therapeutic effects, initial in-vitro data positions UDP-003 as a promising candidate for future therapies.

## 6. Acknowledgements

This work was supported by funding from Cyclarity Therapeutics, National Institute of Aging of the National Institutes of Health under award number 1R41AG071265-01A1, and the National Academy of Medicine under catalyst award number 2000012740.

## 7. Supplementary material

Supplementary data for this article can be found online.

