## Supplemental Data for "Selective Removal of 7KC by a Novel Atherosclerosis Therapeutic Candidate Reverts Foam Cells to a Macrophage-like Phenotype"

| NAME | FORWARD 5’-->3’ | REVERSE 5’-->3’ |
| --- | --- | --- |
| GAPDH | TGAAGCAGGCATCTGAGGG | CGAAGGTGGAAGAGTGGGAG |
| PPARγ | TGTGGGGATAAAGCATCAGGC | CCGGCAGTTAAGATCACACCTAT |
| MIP1α | GAAGAGTCCCTCGATGTGGCTA | CCCTTTTCTGTTCTGCTGACAAG |
| LOX-1 | TTGGCTGAGGTCCTCGACTG | AGGGAACCACCATGGGGAAG |

**Supplementary Section**

**Supplementary Table 1. Primer Sequences.**

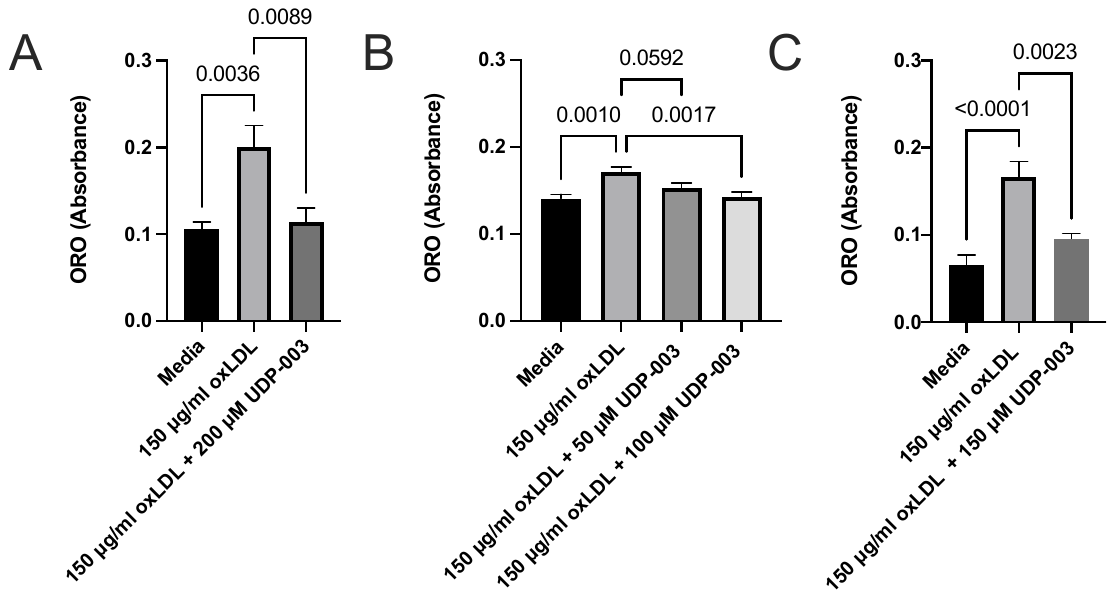

**Supplementary Figure 1. In vitro reduction of intracellular lipid droplets in oxLDL-treated RAW 264.7 macrophages with UDP-003**

Quantification of intracellular lipid droplets collected from RAW 264.7 macrophages was through Oil Red O. Under all modalities, the positive control macrophages were treated with 150 µg/ml oxLDL for 24 h. The macrophages were treated under (**A**) preventative modality, (**B**) sequential modality, and (**C)** prolonged modality. Data is expressed in average absorbance with the baseline absorbance subtracted ± SE bars from at least three independent replicates. Statistical analysis was performed using a one-way analysis of variance (ANOVA).

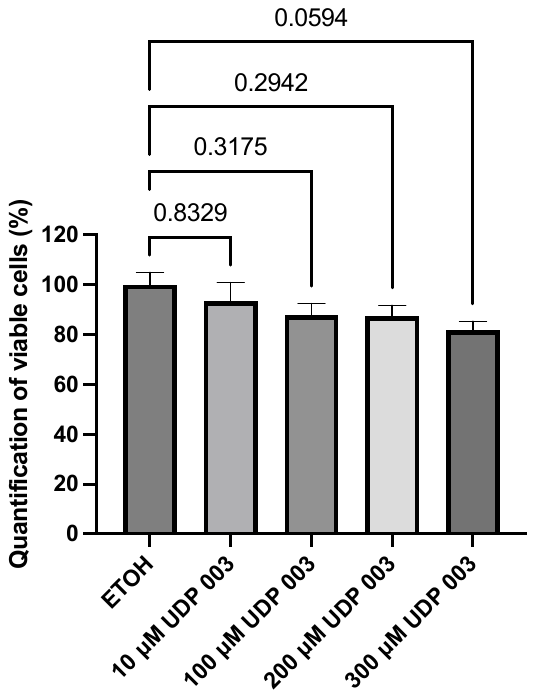

**Supplementary Figure 2. Determination of UDP-003 toxicity of RAW 264.7 macrophages**

RAW 264.7 cells (0.5×10^6^ cells/well in 12-well culture plates) were treated with UDP-003 (10-300 µg/ml) for 24 h. The cell viability was measured using propidium iodide with the BD Accuri C6 flow cytometer with excitation maxima of 535 nm and an emission maxima of 615 nm. The media bar represents negative control without any UDP-003. Data, obtained from triplicate repeats at least, are shown as mean ± SE. Statistical analysis was performed using a one-way analysis of variance (ANOVA).
